# Phage Resistance Mechanisms Increase Colistin Sensitivity in *Acinetobacter baumannii*

**DOI:** 10.1101/2021.07.23.453473

**Authors:** Xiaoqing Wang, Belinda Loh, Yunsong Yu, Xiaoting Hua, Sebastian Leptihn

**Author notes:** Correspondence can be addressed to: Prof. Xiaoting Hua, Sir Run Run Shaw Hospital, Hangzhou, China, East Qingchun Rd 3, Jianggan District, Hangzhou 310016, P. R. China; Prof. Sebastian Leptihn, Infection Medicine, Biomedical Sciences, Edinburgh Medical School, College of Medicine and Veterinary Medicine, The University of Edinburgh, 47 Little France Crescent, EH16 4TJ Edinburgh, United Kingdom.

## Abstract

Few emergency-use antibiotics remain for the treatment of multidrug-resistant bacterial infections. Infections with resistant bacteria are becoming increasingly common. Phage therapy has reemerged as a promising strategy to treat such infections, as microbial viruses are not affected by bacterial resistance to antimicrobial compounds. However, phage therapy is impeded by rapid emergence of phage-resistant bacteria during therapy. In this work, we studied phage-resistance of colistin sensitive and resistant *A. baumannii* strains. Using whole genome sequencing, we determined that phage resistant strains displayed mutations in genes that alter the architecture of the bacterial envelope. In contrast to previous studies where phage-escape mutants showed decreased binding of phages to the bacterial envelope, we obtained several not uninfectable isolates that allowed similar phage adsorption compared to the susceptible strain. When phage-resistant bacteria emerged in the absence of antibiotics, we observed that the colistin resistance levels often decreased, while the antibiotic resistance mechanism *per se* remained unaltered. In particular the two mutated genes that conveyed phage resistance, a putative amylovoran-biosynthesis and a lipo-oligosaccharide (LOS) biosynthesis gene, impact colistin resistance as the mutations increased sensitivity to the antibiotic.

## Introduction

Antimicrobial resistance (AMR) is a global concern. The overuse of antibiotics in human medicine and the misuse of even last resort antimicrobial compounds such as colistin in agriculture, is contributing to the increasing number of antibiotic resistant bacterial pathogens (1). For most antibiotics, molecular mechanisms of resistance have evolved which are quickly distributed throughout bacterial populations by horizontal gene transfer. This is primarily mediated by plasmids and ICEs (2–4). At the same time, the slow and expensive discovery process and clinical development of antimicrobial compounds together with the lack of monetary incentives have resulted in continuously decreasing numbers of effective drugs to treat bacterial infections (5, 6).

The last resort antibiotic colistin is often accompanied by strong side effects, have to be deployed for infections by multidrug resistant pathogens. Here, colistin has gained importance for clinical use. However, colistin resistance in pathogens is increasing as well. Phage therapy has emerged as a promising strategy to treat drug resistant bacterial infections, as viruses are not affected by resistance to antimicrobial compounds (7–9). Phage therapy is the use of lytic phages that have the ability to inactivate pathogens. However, phage-resistance, i.e. the emergence of bacterial mutants that are resistant to a therapeutic phage, is commonly observed (10). Several solutions have been explored in the past, such as combinational phage-antibiotic therapeutic courses, where synergetic effects are often observed, or the deployment of phage mixtures (“phage cocktails”). Yet, in the majority of clinical trials phage-resistance occurs (11). Therefore, it is important to understand the mechanisms that enable bacteria to gain resistance to phages and the consequences of selection. In addition, identifying target molecules that facilitate phage infection and deployment of phages that do not bind the same receptors has been proposed to decrease the likelihood of phage-resistance (12).

In order to understand molecular mechanisms of phage-resistance, we investigated a phage-pathogen system consisting of the type strain of *A. baumannii* and a colistin-resistant mutant of ATCC17978. We employed whole genome sequencing of phage-escape mutants that emerged after co-incubating the novel phage Phab24 that infects both strains. We found that genes abolishing infection are primarily involved in the biogenesis of the envelope of the bacterial host, namely LPS (LOS) and capsular polysaccharides. The two genes we identified in *A. baumannii* are involved in the biosynthesis of the bacterial envelope; the putative amylovoran biosynthesis gene *amsE* is involved in capsule formation while *lpsBSP* plays a role in lipo-oligosaccharide (LOS) biosynthesis. While an *amsE* deletion leads to decreased adsorption, the mutation in the *lpsBSP* gene does not alter binding of the phage to the bacterial surface. Gene engineered strains introducing the observed mutations one at a time in the parental strain, and their complementation with the wildtype gene *in trans*, confirmed the role of the genes in phage-resistance. *In vitro* evolution experiments resulted in the selection of escape mutants with decreased antibiotic resistance, with mutations in *amsE* contributing to this effect.

## Results

### Isolation of Phage Phab24 that infects Colistin-Resistant *A. baumannii* Strain XH198

Colistin resistance is mediated by a fundamental change in the bacterial membrane composition. A mutation in gene *pmrB* (G315D) results in surface modification of the bacterial envelope, preventing efficient colistin binding and thus mediating resistance to the antibiotic (13–17). To study phage-resistance of bacteria *in vitro,* we used the colistin resistant *Acinetobacter baumannii* strain XH198 and its non-resistant parental strain ATCC17978. Strain XH198 was obtained by an *in vitro* evolution experiment and exhibits altered LOS molecules, which has also been observed in clinical isolates, rendering colistin ineffective (14). We isolated several phages that are able to infect both, XH198 and ATCC17978. To focus on one particular virus-host system, we selected a phage that we named Phab24, for <u>Ph</u>age *<u>A</u>cinetobacter <u>b</u>aumannii* number <u>24</u> (genome accession number: MZ477002) (Figure 1A). Whole genome analysis using the programme PhageAI (https://phage.ai/) revealed that bacteriophage Phab24 is virulent (lytic) with a 93.76% confidence in prediction (18).

**Figure 1:**
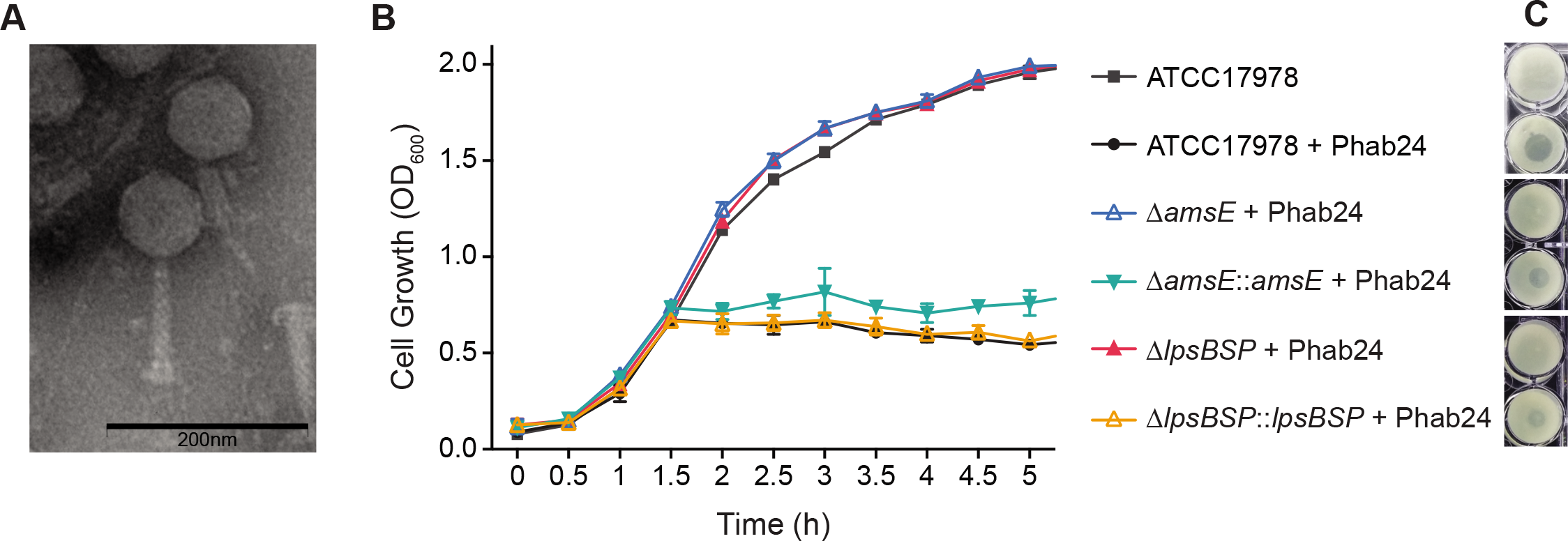
Phage Phab24 and effect on *A. baumannii* strain growth dynamics. (A) Transmission electron micrograph of Phab24 (negative staining). Phage Phab24 belongs to Myoviridae which have contractible tails (as seen on the right). Bar: 200nm. (B) Growth curves of the *A. baumannii* reference strain ATCC17978 and phage-resistant reconstructed strains with introduced genomic mutations in the putative amylovoran biosynthesis gene *amsE* or in the putative LPS biogenesis gene (“*lpsBSP*”), and their plasmid-complementations (::*asmE* and ::*lpsBSP*) in the presence or absence of phage Phab24. (C) Spot testing of Phab24 on agar containing phage-resistant and susceptible strains.

### Isolation and characterisation of phage-resistant bacterial mutants

We first used the phage-host system of Phab24 and ATCC17978, to study the emergence of phage-resistant bacterial mutants, by co-incubating both in liquid media. Subsequently, bacteria were plated on solid media from which we randomly picked 80 bacterial colonies (R1-R80). Surprisingly, two isolates displayed susceptibility to Phab24, possibly persister cells (19) while the remaining 78 were resistant to Phab24. Similarly, from co-incubating Phab24 with XH198, the colistin-resistant derivative of the ATCC reference strain, we isolated 400 colonies at random. Next, we sequenced the whole genome of six of the phage-resistant isolates of ATCC17978 (R5, R10, R22, R23, R39, R70) and nine of XH198 (R81, R83, R86, R115, R125, R130, R132, R134, R137). Using Breseq (20), we found several point mutations as well as deletions or insertions in genes that might mediate phage-resistance, which were confirmed by PCR and subsequent sequencing (Table 1 and Supplemental table 1). The most common gene to have mutations codes for a putative LPS biosynthesis protein, which we subsequently called *lpsBSP* (gene AUO97_03485). Here, we observed a single nucleotide deletion or an IS insertion, indicating parallel emergence of phage resistance. We also found mutations in genes predicted to code for glycosyltransferases (glycosyl transferase 1, AUO97_06920 and glycosyl transferase 2 AUO97_03485), phosphohydrolase *phoH* and the putative ABC transporter *abcT*. While we found many mutations in genes that were later identified to be irrelevant for phage-resistance, we also identified two types of mutations (a frameshift or an insertion) in a gene called *amsE* (AUO97_06900), which encodes a putative amylovoran biosynthesis protein.

**Table 1:** Resistant strains and mutations conveying phage resistance and outcome of wildtype gene complementations *in trans*. Mutations were either detected by whole genome sequencing (strains R5, R10, R22, R23, R39, R70, R81, R83, R86, R115, R125, R130, R132, R134, R137), or screened for using mutation-specific primers.

| Resistant Isolate | Mutation | Putative gene function | Complementation |  |
| --- | --- | --- | --- | --- |
|  |  |  | Yes | No |
| R1 | *lpsBSP*<br>(IS4 family insertion) | LPS/ LOS biosynthesis | *lpsBSP* |  |
| R2 | *lpsBSP*<br>( 535<sup>th</sup> A loss, p.Asn179Ile fs Ter7) | LPS/ LOS biosynthesis | *lpsBSP* |  |
| R5 | *amsE*<br>(IS4 insertion) | Amylovoran biosynthesis |  | *amsE* |
|  | *lpsBSP*<br>( 535<sup>th</sup> A loss, p.Asn179Ile fs Ter7) | LPS/ LOS biosynthesis |  | *lpsBSP* |
| R7 | *amsE*<br>(IS5 insertion) | Amylovoran biosynthesis | *amsE* |  |
| R8 | *lpsBSP*<br>(535<sup>th</sup> A loss, p.Asn179Ile fs Ter7) | LPS/ LOS biosynthesis | *lpsBSP* |  |
| R10 | *amsE*<br>(559<sup>th</sup> T loss, p. Leu187Tyr fs Ter3) | Amylovoran biosynthesis | *amsE* |  |
| R12 | *lpsBSP*<br>(535<sup>th</sup> A loss, p.Asn179Ile fs Ter7) | LPS/ LOS biosynthesis | *lpsBSP* |  |
| R13 | No PCR product of *amsE* | Amylovoran biosynthesis |  | *amsE* |
| R28 | *amsE*<br>(transposase insertion) | Amylovoran biosynthesis |  | *amsE* |
| R33 | *amsE*<br>(IS5 insertion) | Amylovoran biosynthesis |  | *amsE* |
| R35 | *lpsBSP*<br>(535<sup>th</sup> A loss, p. Asn179Ile fs Ter7) | LPS/ LOS biosynthesis | *lpsBSP* |  |
| R39 | *lpsBSP*<br>(535<sup>th</sup> A loss, p.Asn179Ile fs Ter7) | LPS/ LOS biosynthesis | *lpsBSP* |  |
| R54 | *amsE* (transposase insertion) | Amylovoran biosynthesis |  | *amsE* |
| R78 | *lpsBSP*<br>(IS5 family insertion) | LPS/ LOS biosynthesis | *lpsBSP* |  |
|  |  |  | Yes | No |
| *amsE* KO | Markerless knock out the entire *amsE* on the background of ATCC17978 | Amylovoran biosynthesis | *amsE* |  |
| *lpsBSP*-delta A | Substitute for the wild type *lpsBSP* on the background of ATCC17978 | LPS/ LOS biosynthesis | *lpsBSP* |  |
| R81 | *amsE* (transposase insertion) | Amylovoran biosynthesis | *amsE* |  |
| R83 | No PCR product of *amsE* | Amylovoran biosynthesis |  | *amsE* |
| R86 | *amsE* (transposase insertion) | Amylovoran biosynthesis | *amsE* |  |
| R115 | *amsE* (transposase insertion) | Amylovoran biosynthesis | *amsE* |  |
| R130 | *amsE* (transposase insertion) | Amylovoran biosynthesis | *amsE* |  |
| R132 | *amsE* (transposase insertion) | Amylovoran biosynthesis |  | *amsE* |
| R134 | *amsE* (transposase insertion) | Amylovoran biosynthesis | *amsE* |  |
| XH198<br>*amsE* KO | Markerless knock out the entire *amsE* on the background of XH198 | Amylovoran biosynthesis | *amsE* |  |
| XH198<br>*lpsBSP*-KO | Markerless knock out the entire *lpsBSP* on the background of XH198 | LPS/ LOS biosynthesis | *lpsBSP* |  |
| R518 | *lpsBSP*<br>(535<sup>th</sup> A loss, p.Asn179Ile fs Ter7) | LPS/ LOS biosynthesis | *lpsBSP* |  |
| R587 | *lpsBSP*<br>(535<sup>th</sup> A loss, p.Asn179Ile fs Ter7) | LPS/ LOS biosynthesis | *lpsBSP* |  |

Yes: After the transformation of the corresponding plasmid which contains the WT gene, the R variant can get infected by Phab24.
No: After the transformation of the corresponding plasmid which contains the WT gene, the R variant cannot get infected by Phab24.
R1-R80 are Phab24 escape mutants from ATCC17978.
R81-R587 are Phab24 escape variants from XH198.
Based on the NCBI accession number CP018664.1, ATCC17978:
AUO97-06900, *amsE*, 828nt
AUO97-03485, *lpsBSP*, 768nt

To demonstrate that the mutations indeed render a strain non-susceptible, we constructed several plasmids encoding the wildtype genes in the *A. baumannii* shuttle vector pYMAb2 (21, 22). We then introduced the episomal elements into the phage-resistant (R) -mutants. The expression of the wildtype genes of the LPS biosynthesis protein *lpsBSP* (e.g. in R1) as well as of *amsE* (e.g. in R7) did restore phage sensitivity, unless they were both present in the strain (e.g. strain R5) (Table 1). Some phage-resistant isolates could not be of complemented by wildtype genes coding for some of the mutations we observed (e.g. the membrane transport protein abcT, phoH, or actP), indicating the presence of additional, as yet unidentified, mutations that confer resistance (Supplemental table 1).

Due to the obvious complexity of the resistance mechanisms, we decided to focus on *lpsBSP* and *amsE*. To demonstrate the role of these genes in phage susceptibility, in addition to complementations *in trans*, we engineered the mutations into the parental *A. baumannii* strain (ATCC17978), or created knock-out mutants of genes. When mutations are introduced into the gene coding for *lpsBSP*, the bacterium is rendered “immune” to phage Phab24 (Figure 1B). Similarly, a reconstructed mutant in which the *amsE* gene is deleted cannot be infected by the phage, clearly demonstrating the role of these two genes in phage susceptibility. Both reconstructed mutants can be infected by Phab24, when the complementing wildtype genes are expressed *in trans*, from plasmids (Figure 1A and B).

### Attachment of Phab24 to the surface of phage-resistant mutants

As the mutations found in *lpsBSP* and *amsE* code for putative proteins involved in LOS and capsule formation, respectively, we concluded that the surface of the phage-resistant mutants may exhibit modifications compared to the wildtype. Changes on the surface of mutant strains might therefore lead to a reduction in binding of phages to bacteria. We therefore performed binding assays to assess the quantity of phages that bind to the bacterial envelope. To this end, we co-incubated Phab24 with the phage-resistant mutants and controls for 20 minutes and subsequently determined the phage titre left in the supernatant (Figure 2). The positive control, strain ATCC17978, was able to reduce the titre by a factor of more than 1000-fold, while the negative controls (no bacteria or *A. baumannii* strain XH194 resistant to Phab24 infection) resulted only in a minor reduction in the number of phage particles in the supernatant. Interestingly, Phab24 bound ATCC17978 with an apparent binding affinity higher than XH198. The *amsE* knockout on the backbone of ATCC17978 showed reduced bacteriophage binding, similar to the negative controls. However, upon complementation with the wildtype *amsE* gene *in trans*, binding of Phab24 was restored to a level comparable to parental ATCC17978. However, the *IpsBSP* knockout and its derivative complemented with wildtype the *lpsBSP* knockout *lpsBSP* both bound Phab24 avidly just like the parental ATCC17978 strain. Surprisingly, with the exception of *amsE* mutants, phage Phab24 still binds to most phage-resistant strains (Figure 2B).

**Figure 2:**
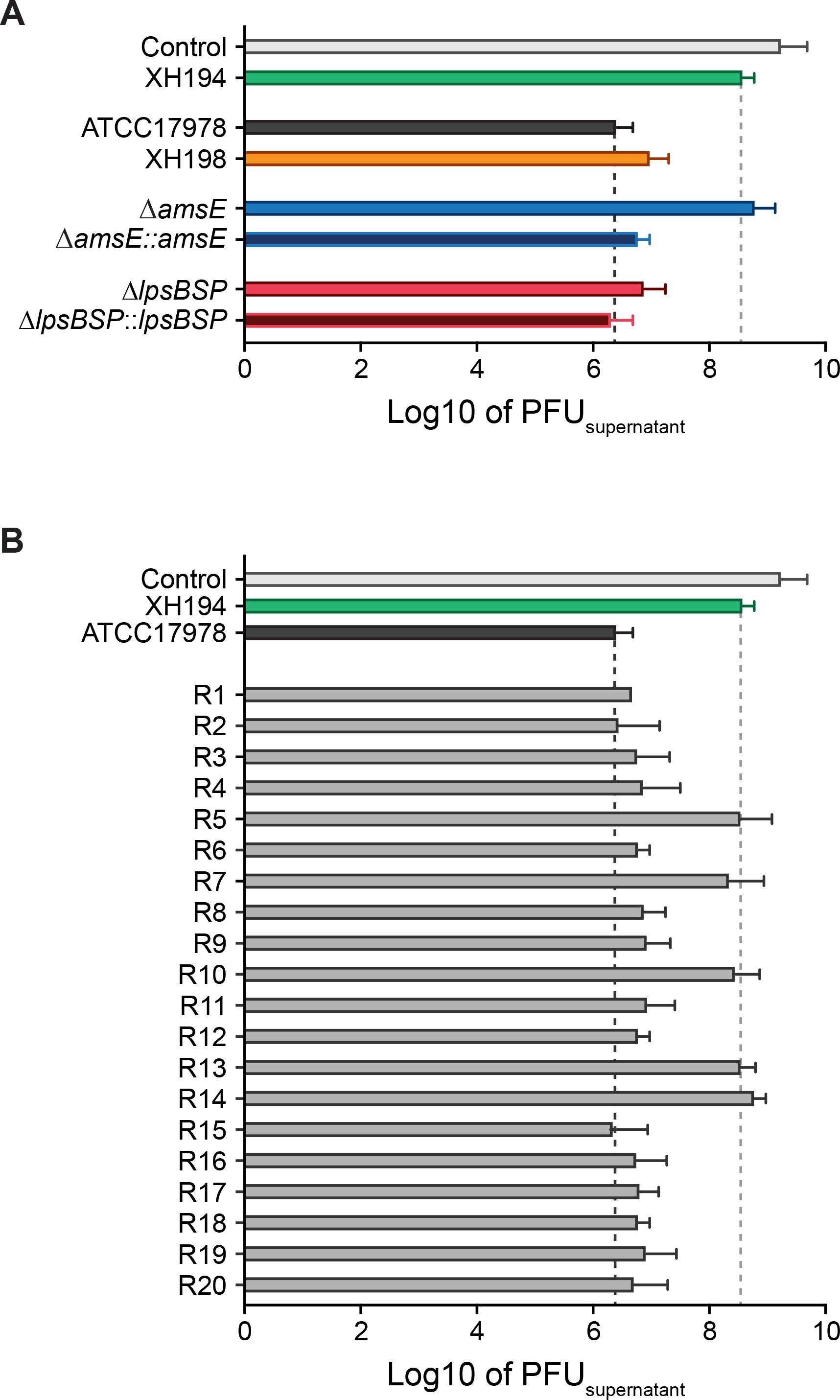
Attachment assay of Phab24 to bacteria. Titre of free phages detected in media after incubation with (A) reference strain ATCC17978, colistin resistant derivate XH198 as well as the *amsE* and *lpsBSP* genetically engineered strains (Δ) and their complementations (::), and (B) Phab24 phage-resistant colonies, “R”, derived from ATCC17978. Control: phage Phab24 incubated without bacteria. XH194: bacterial strain that is colistin resistant and is not infected by Phab24.

### The bacterial envelope is altered in both, the LPS biosynthesis protein mutant and the *amsE* mutant

LPS often serves as a co-receptor in phage binding. As a disruption in the gene coding for a LPS biosynthesis protein (*lpsBSP*) was observed, we determined if the mutations in the isolated strains lead to a change in LPS composition. Mass spectrometry of isolated lipid A, obtained using the hot aqueous phenol extraction method, was performed to compare the reconstructed mutants (on the ATCC17978 backbone) with the ATCC17978 control (Figure 3A and B). In addition, we analysed the samples of the plasmid-complemented strain that allows the expression of *lpsBSP in trans*. Previously, it was established that the m/z of *A. baumannii* lipid A is featured as a prominent peak at 1,910, which was identified as a singly deprotonated lipid A structure that contains two phosphate groups and seven acyl chains (i.e., diphosphoryl hepta-acylated lipid A) (23). In our experiments, this peak was observed in all samples (Figure 3). While the mass spectrum of the KO strain (Figure 3B) shows molecules with mass / charge (m/z) values of 1,910 and smaller, the reference spectrum (Figure 3A) exhibits several additional small peaks larger than 2,000. These higher molecular weight peaks are more prominent in the plasmid-complemented strain (Figure 3C). While the identification of molecules that lead to the occurrence of these peaks with higher m/z values is still outstanding, they possibly represent modified Lipid A molecules. As these peaks are indistinguishable from the baseline spectrum recorded with the KO strain sample (Figure 3B), it may be reasonable to conclude that the disruption in *lpsBSP* results in a modification in the bacterial surface structure. The gene disruption might ultimately prevent the incorporation of one or more types of modified LPS molecules into the bacterial envelope which are able to facilitate specific Phab24 phage binding (15, 23).

**Figure 3:**
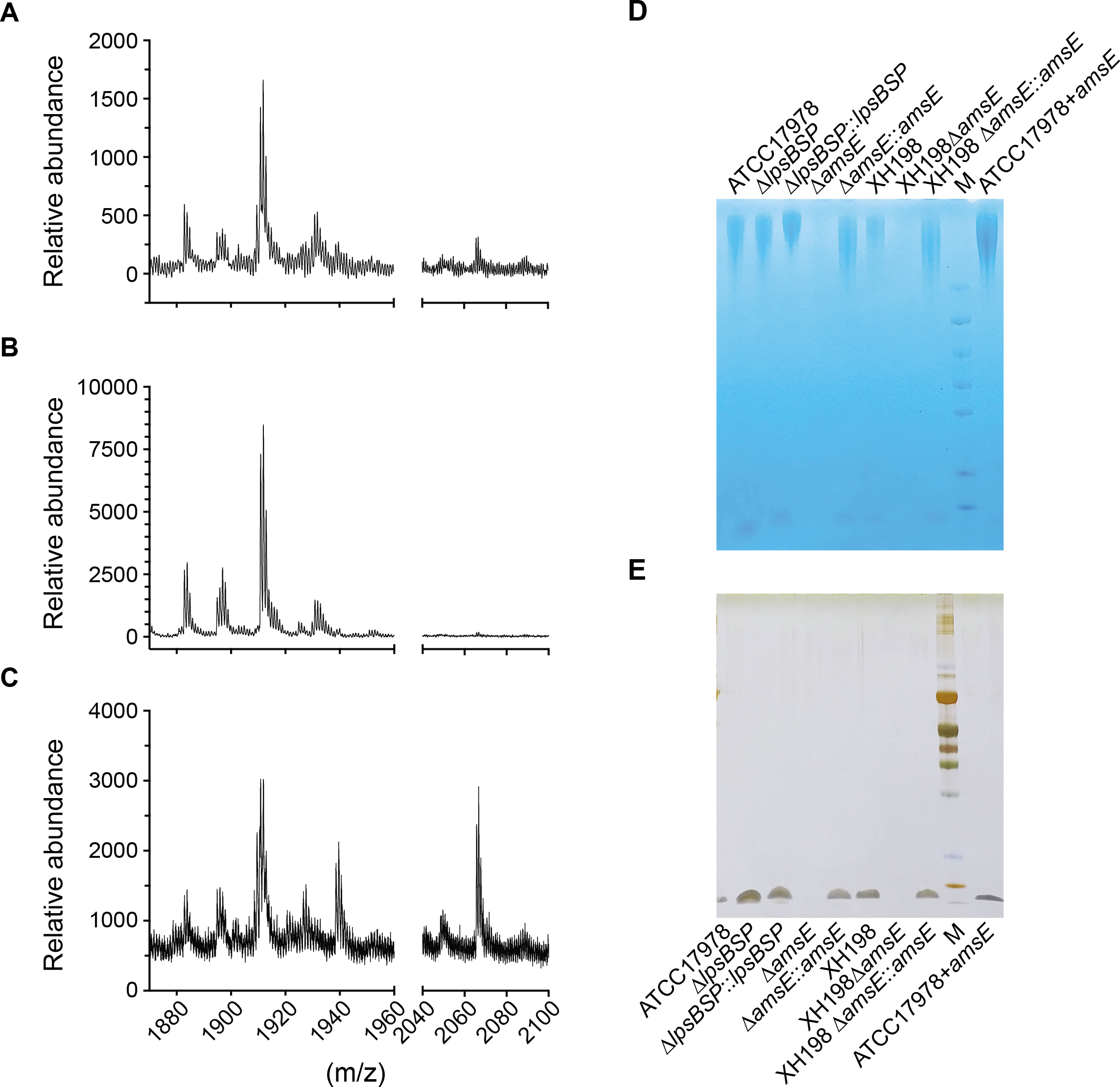
Composition of the bacterial envelope. (A to C) Mass spectrometry (positive mode) analysis of lipid A isolated using aqueous phenol extraction from (A) Wildtype strain ATCC17978, (B) Phab24 resistant, genetically introduced reconstructed *lpsBSP* gene mutant on ATCC17978, (C) Plasmid complemented Phab24 resistant strain. (D&E) SDS-PAGE gel of isolated capsular polysaccharides stained with (D) Alcian blue, which allows the detection of acidic polysaccharides, and (E) silver staining, which detects lipids and proteins/peptides.

As *amsE* is a putative amylovoran biosynthesis protein involved in the production of amylovoran, an acidic exopolysaccharide (EPS), we isolated the oligosaccharides of the reconstructed *amsE* mutant and to compare strain ATCC17978, and again performed mass spectrometry without the intension to characterise the molecules observed in the spectra. We only used the method to compare the mass spectrometry results to each other. Perhaps unexpectedly, we did not observe any additional presence or absence of peaks in the spectra across 180-3200 m/z (Supplemental Fig. 1, 2, 3). While ratio and intensity varied for some peaks, there is no indication of the absence of specific polysaccharides, or the presence of others, in the *amsE* mutant. The relative quantity of the polysaccharides cannot be firmly established with the method we employed, and thus we attempted to determine if differences can be seen on SDS PAGE gels which allow a separation of molecules based on size, while also allowing a quantitative analysis. Here, we observed that the genetically engineered *amsE* mutants of ATCC17978 or of XH198 exhibited a massive reduction in material on the gel compared to the complemented strains (and the reference strains) where *amsE* was expressed *in trans* (Figure 3D, E). A gel with Alcian blue, which stains acidic polysaccharides, shows that the *amsE* mutants of both, ATCC17978 and XH198, contain almost undetectable amounts of material, while the plasmid-complemented strains are similar to the wildtype level. In addition to large molecular weight bands, we also observe small molecule components which are stained by Alcian blue, but also by silver ions. As silver staining allows the sensitive detection of lipids and proteins but not of the saccharides (and polypeptides can be excluded due to the preparation which includes a Proteinase K digestion step), the smaller molecules possibly indicate the presence of lipid-saccharide conjugates in the samples, which again, are absent in the *amsE* mutants. Although the mass spectrometry data did not indicate any changes in saccharide composition, the quantitative method of size-separated oligosaccharides on gels indicate that the bacterial envelope surface (possibly not the composition) of the *amsE* mutant is clearly different from that of the *A. baumannii* reference strain.

### Changes in cell morphology and capsule formation in phage-escape mutants

On the molecular level we could confirm that the composition of the cell envelope is different in the *lpsBSP* mutant while we found no indication for a qualitative change in the *amsE* mutant, although the mutant appears to produce substantially less exopolysaccharides. However, it is difficult to conclude that the surface structure of the cells is altered by the mutations. We first employed Transmission electron microscopy on thin sections of resin-embedded cells with inconclusive results (Supplemental Figure 2). We then used Scanning Electron Microscopy where, both, ATCC17978 and XH198 cells, appear similarly smooth and rod-shaped. A similar morphology could be observed in the case of cells containing the *lpsBSP* mutant; when complemented *in trans* by the functional gene, a slightly more “shrivelled” -possibly desiccated- structure was observed. A very even, smooth surface was seen when the *amsE* mutant was complemented with the functional gene *in trans* while the surface of the *amsE* mutant itself appeared less smooth. In addition, the cells of the *amsE* mutant appear more rounded and adherent to each other, forming clusters (Figure 4). We also observed the formation of mucoid-like strings in preparations of the *amsE* mutant which were absent in all other samples (Supplemental Figure 3).

**Figure 4:**
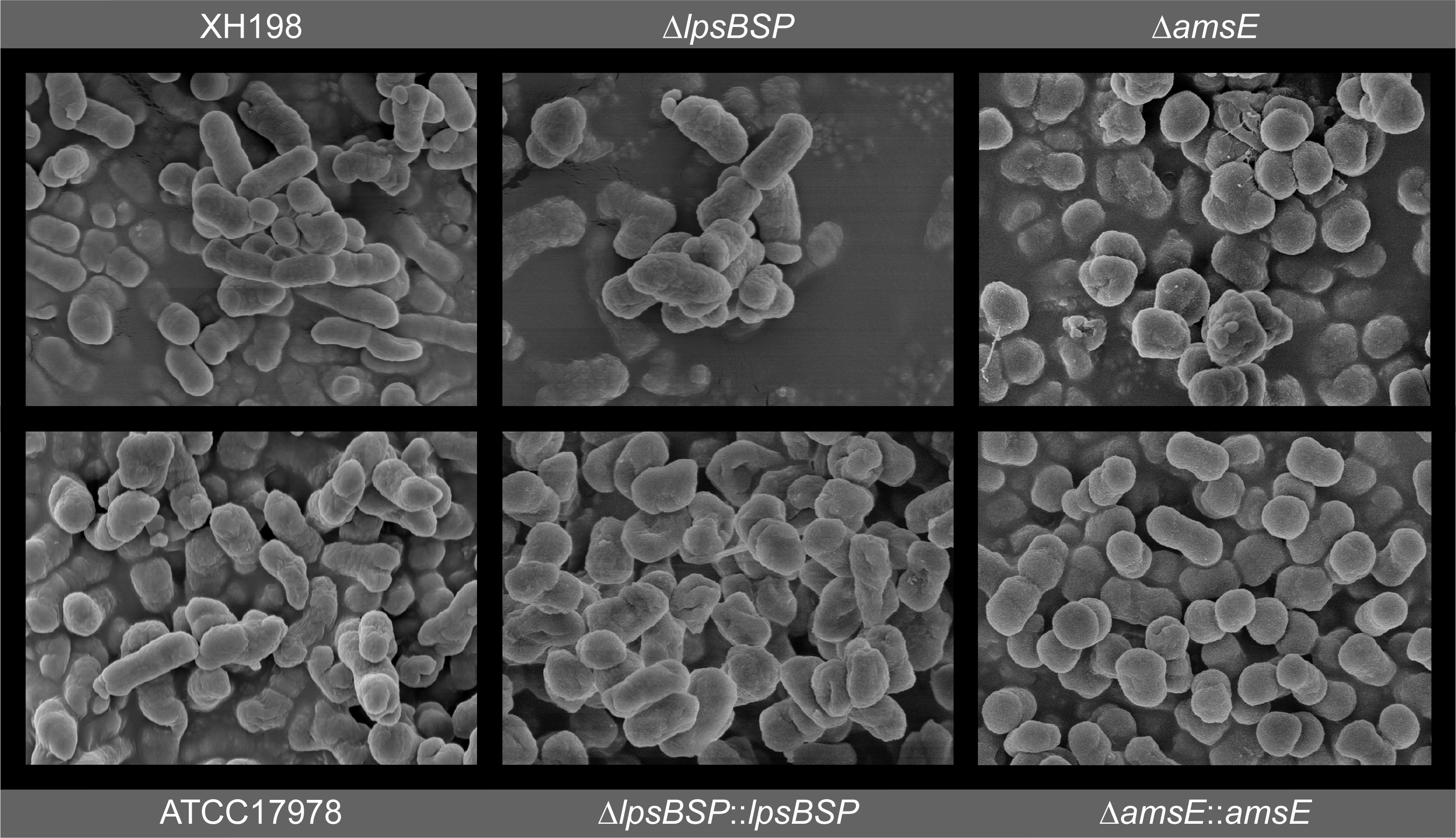
Surface structure of bacterial cell envelope. Scanning Electron Microscopy. *ΔlpsBSP*: knock-out of *lpsBSP* gene in the ATCC17978. *ΔlpsBSP::lpsBSP*: knock-out of *lpsBSP* gene which was complemented with the wildtype *lpsBSP* gene *in trans*. *ΔamsE*: knock-out of *amsE* gene in the ATCC17978. *ΔamsE::amsE*: knock-out of *amsE* gene which was complemented with the wildtype *amsE* gene *in trans*.

The observed clustering and appearance of “slime” on the surface of the *amsE* mutant may be caused by exopolysaccharide and/or biofilm formation. We therefore assessed the production of material produced by cells grown for three days to retain the dye crystal violet, often used to quantify the extent of biofilm produced by bacteria. Here, we observed that biofilm formation (the ability to retain the dye) was most pronounced in the colistin resistant strain XH198, but remained low in all other strains (Figure 5). Compared to the reference strain ATCC17978, the *amsE* mutant showed reduced biofilm formation (∼42 % reduction compared to ATCC17978), while plasmid complementation led to a slight increase in biofilm (∼117% of that of ATCC17978) (Figure 5A). In the case of the *lpsBSP* mutant, no significant difference in biofilm formation of the complemented or non-complemented strains was observed. Thus, the “mucoid” appearance and aggregation observed in SEM preparations of the cells in the *amsE* mutant cannot be explained by the formation of biofilms. Interestingly, we observed the formation of highly fragile membraneous structures in the multi-well plates for this mutant that, however, did not retain the dye, and were washed away easily (Supplemental Figure 4). In addition to the formation of biofilms we also employed crystal violet staining of capsules of planktonic cells. Biofilms are usually formed if bacteria are incubated for a long duration and allowed to sediment and form clusters. If cells are incubated for shorter times under shaking, biofilm formation does not occur. It is reasonable to assume that the amount of dye that is retained by the capsule correlates with the quantity material present for the dye to embed in (i.e. more dye is retained by more extensive capsules) and/ or the density of the packing of the capsule material (i.e. released quicker if the material is less compact). The ATCC17978 strain retained substantially more dye compared to the *amsE* mutant while a complementation results in the same levels observed for the reference strain (Figure 5B). One interpretation is that the *amsE* mutant strain can absorb less dye due to a smaller capsule which would correlate with the results of the capsular material separated on SDS gels (Figure 3). The *lpsBSP* mutant exhibits slightly more absorption of the dye, while the complemented strain shows levels identical to ATCC17978.

**Figure 5:**
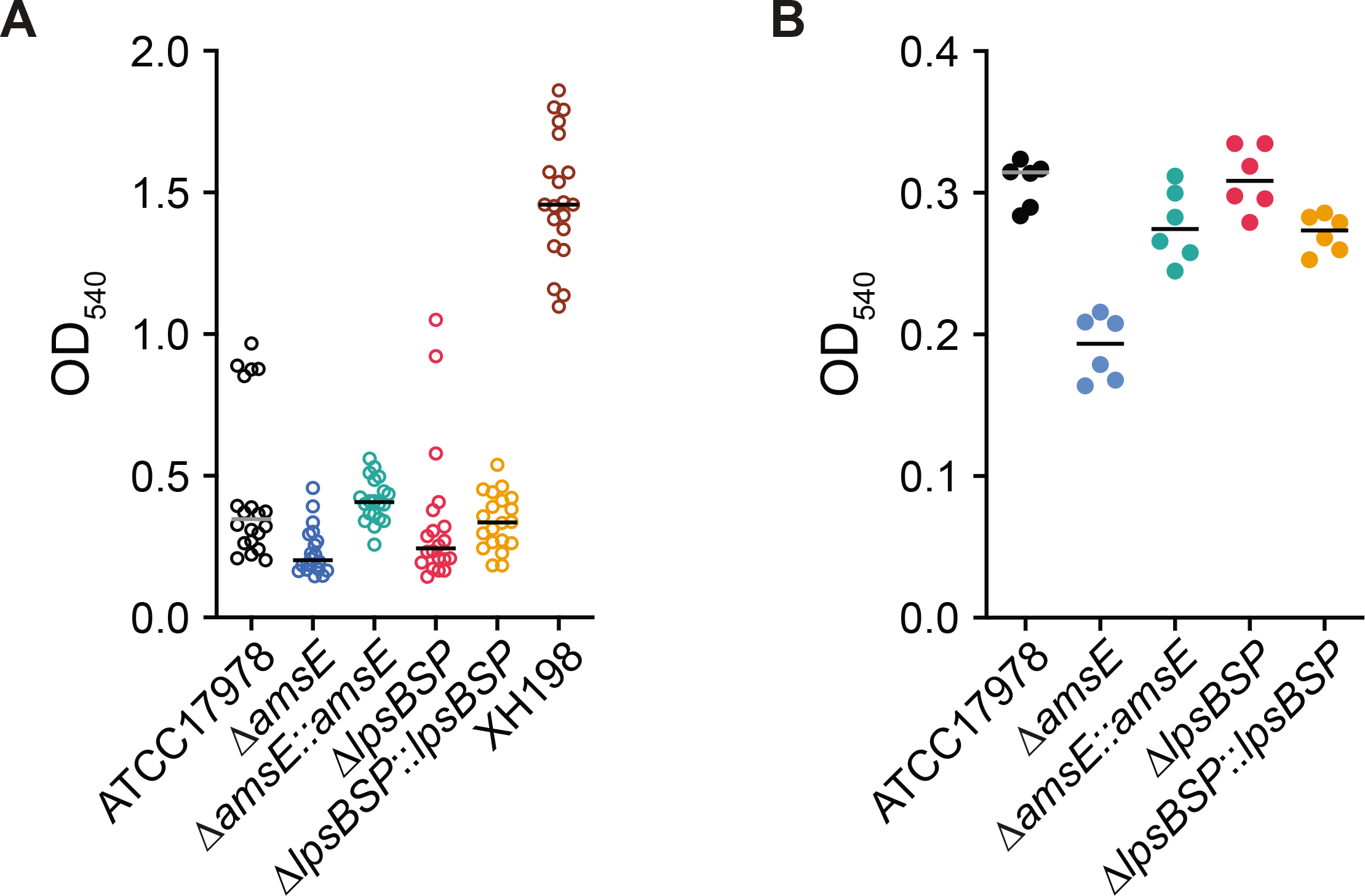
Biofilm formation and capsular stain. (A) Biofilm formation assessed by the ability to retain crystal violet (CV) dye. Strains were incubated for 72 hours in multi well plates without shaking. (B) Capsule stain of planktonic cells incubated for 8 hours with shaking. Cells were stained with CV to determine the ability of the bacterial capsule to retain the dye. Ethanol was used to destain the capsule and release the CV dye which was then detected by measuring absorbance at 540nm.

### AmsE and LPS biosynthesis mutants are less virulent *in vivo*

As part of the extended bacterial surface structure, to which capsules can be included, LPS/LOS and capsular molecules often contribute to virulence of strains (24–27). We therefore investigated how the genes that conferred phage-resistance impact the virulence of the phage-escape mutants. To this end, we used the insect larva model *Galleria melonella*, to assess the virulence of the reference strains (ATCC17978, XH198), compared to the single gene mutant strains (*amsE*, *lpsBSP*), and those complemented *in trans*. The mutation in *lpsBSP* impacts the virulence of the strain only to a small extent, with slightly increased survival rates compared to ATCC17978 (Figure 6). In contrast, the *amsE* KO strain had a significantly reduced virulence, demonstrating the importance of *amsE* on the pathogenicity of *A. baumannii*. When the *amsE* mutant strain was complemented with a plasmid expressing the wildtype gene under a constitutive promotor, the virulence was significantly increased compared to ATCC17978, possibly due to a higher expression level. Similarly, albeit to a lesser extent, virulence of the strain with the *lpsBSP* mutation increased when the wildtype gene was expressed *in trans*.

**Figure 6:**
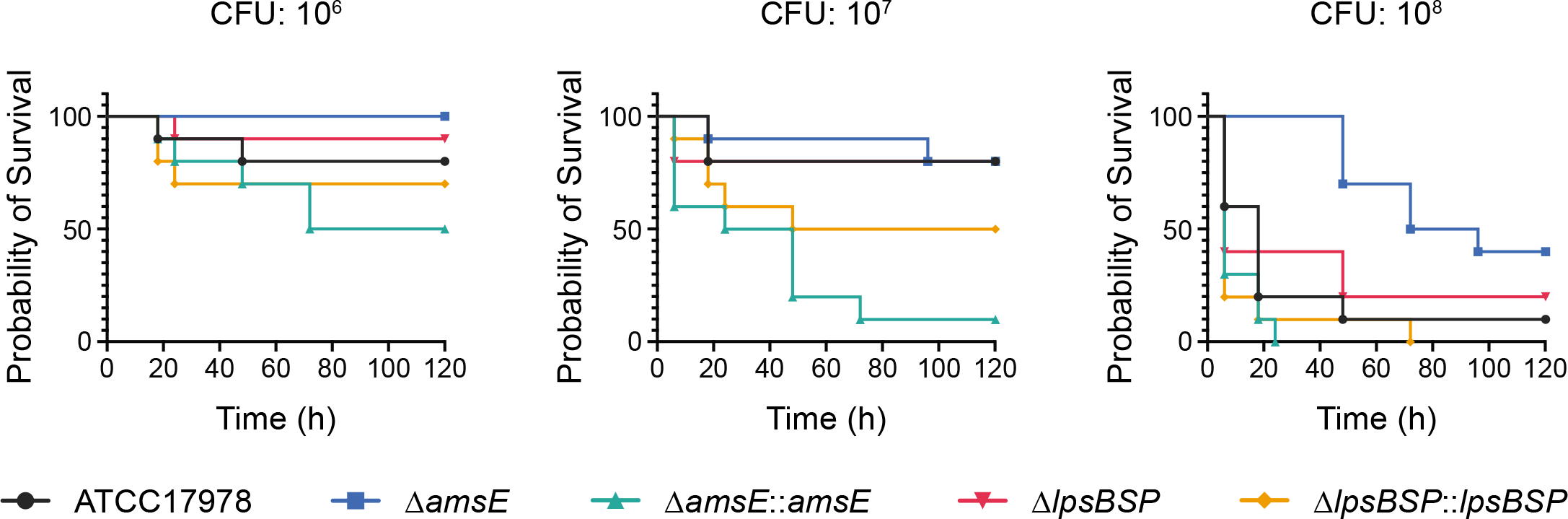
*In vivo* virulence tests of *A. baumannii* strains and the Phab24 resistant isolates. Survival of *Galleria melonella* larvae over 120 h after injection with (A) 10^6^ colony forming units (CFU), (B) 10^7^ CFU, and (C) 10^8^ CFU of *A. baumannii* strains. Each group consisted of 10 larvae. Shown is a representative experiment of 4 independent repeats.

### Phage-resistance mutations in *amsE* decrease colistin resistance

Specific alterations in membrane composition are the molecular basis for colistin resistance. We found that phage Phab24 binds to surface exposed molecules of ATCC17978 and XH198, and resistance is mediated by altered envelope structures. To answer the question if the resistance mechanisms have impact on each other, we performed experiments testing the combination of Phab24 together with colistin. During phage therapy, antibiotics are often used in combination with therapeutic phages, as synergistic effects of phage-antibiotic combinations have often been observed (28, 29). In our case, increasing concentrations of phage reduced the apparent MIC of colistin (Figure 7A). One explanation is that phage-resistant mutants show a higher sensitivity to colistin. To address this hypothesis, we investigated if phage-resistant bacteria show higher sensitivity to colistin, testing colony survival of XH198 cultures grown in the absence of colistin but with Phab24 (MOI of 1) (Figure 7B and C). The numbers of colony forming units (CFUs) were subsequently determined on media with different amounts of colistin. Interestingly, we observed that the number of CFU decreases with increasing colistin concentration, regardless whether or not the bacteria were co-incubated with Phab24. Compared to the number of colonies that grew on plates without colistin, the ratio of bacterial colonies dropped by ∼1/4 to ∼3/4 when colistin was present. However, in the presence of Phab24, the ratio of CFUs was reduced to less than 0.01% compared to the count when colistin was absent. This observation might indicate that the selection for phage-resistance leads to the mutations which in turn increase the sensitivity to colistin. To investigate this finding in more detail, we tested the MIC of individual phage-resistant XH198 mutants (Figure 7D). Resistance levels varied widely from 64 to 0.5 mg/L (Supplemental Table 2). The correlation between the observed reduction in colistin resistance with phage resistance is far from trivial. Colistin resistance in the parental strain XH198 is mediated by a mutation in gene *pmrB* (G315D), which remained unchanged (data not shown). Therefore, the molecular basis for increased sensitivity possibly lies in other mutations found in the strains, including those in *amsE* or *lpsBSP*. Thus, we tested the impact of *amsE* and *lpsBSP* on the colistin sensitivity.

**Figure 7:**
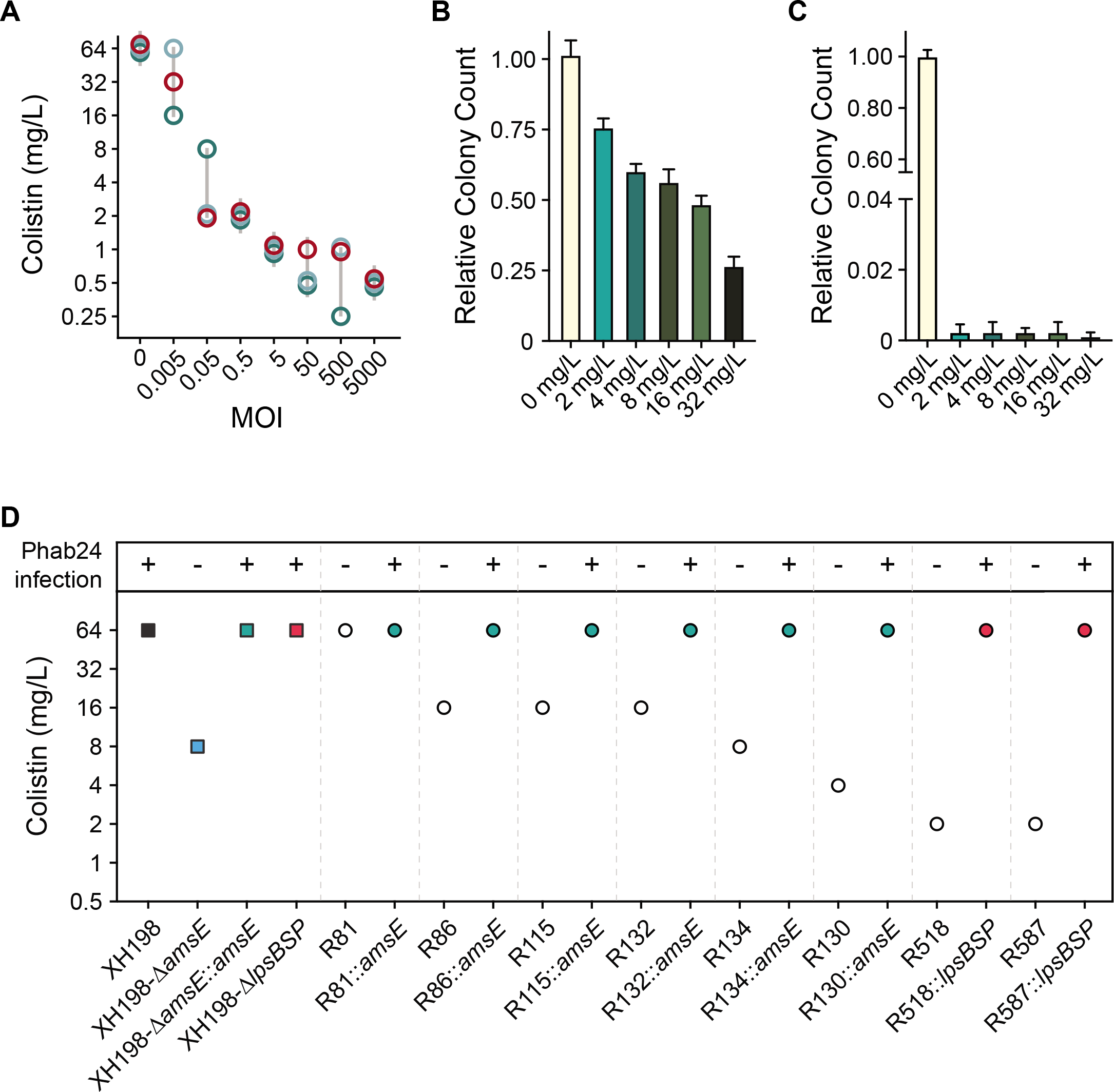
Colistin and Phage resistance. (A) Synergy of colistin with/without phage Phab24. Co-incubation of different numbers of Phab24 (MOI) at different concentrations of colistin (mg/L) at constant cell numbers. Each circle represents an independent experiment. (B&C) Colistin-resistance levels of emerged phage-resistant colonies. Colony count of XH198 incubated for 3 h in the absence of colistin or on colistin plates with different antibiotic concentrations (B) in the absence of phage, and (C) in the presence of Phab24. (D) Colistin MIC of selected Phab24 resistant isolates. Top panel shows if the strain can be infected by Phab24, indicating successful complementation of a mutated gene. The control, strain XH198, exhibits a MIC of 64 mg/L. Phage-resistant isolates displayed reduced levels of resistance varying from 16 - 1 mg/L, except for one strain (R81). The reconstructed *amsE* mutant shows an increased sensitivity to colistin compared to the reference strain XH198, while plasmid complementation with the wildtype gene fully restores the high level of resistance. The genetic deletion of the entire *lpsBSP* gene in XH198 appears does not result in a reduction in colistin resistance. However, strains with mutations that lead to a truncated *lpsBSP* gene product increase colistin sensitivity by 32-fold (R518, R587).

We first tested XH198-derived phage resistant isolates with *lpsBSP* mutations, R518 and R587, where we observed a colistin MIC of 2 mg/ L. Both strains can be complemented with a plasmid-encoded *lspBSP* which then allows infection by Phab24 increasing the MIC to 64 mg/ L (Figure 7D). While the strains may have additional mutations, this might be an indication that the *lpsBSP* mutation increases colistin sensitivity. Curiously, the genetic deletion of the entire *lpsBSP* gene in XH198 (a knock-out) appears does not result in a reduction in colistin resistance. We then tested the reconstructed *amsE* mutant on the backbone of XH198, which showed strongly increased sensitivity to colistin, from 64 mg/ L to 8 mg/ L (Figure 7D). The complementation *in trans* using a plasmid-encoded wildtype *amsE* restored the high resistance level to colistin. Similarly, mutants containing *amsE* mutations exhibiting various degrees of colistin sensitivity, displayed the high level of resistance when complemented e.g. R130 from 4 mg/ L (without plasmid), to 64 mg/ L (with plasmid).

## Discussion

Bacterial phage susceptibility rests on several mechanisms, one of which is binding to receptors and co-receptors. In our work, we have established the role of two genes in permitting the infection of *A. baumannii* strain ATCC17978 and its colistin-resistant derivative, XH198 by phage Phab24. We provide evidence that the genes encode proteins that are involved in the biogenesis of the bacterial envelope and function as phage receptors (or receptor and co-receptor). One gene has a putative function in the biogenesis of LPS (or LOS), which we named *lpsBSP*. The second gene, *amsE*, is likely involved in the biosynthesis of amylovoran for capsular exopolysaccharide production. Previous work has shown that capsular molecules can serve as phage receptors, which we also show describe in this study (30–33). Phage-resistant isolates that displayed mutations in *amsE* did not permit efficient binding or infection by Phab24. These mutations may either lead to the production of altered surface receptors to which the phage cannot bind, or perhaps abolish the production of the “correct” receptor. Surprisingly, we found that binding is unaltered in strains displaying mutations in *lpsBSP*, yet infection does not occur, which contrasts with previous reports (34, 35). For some phages, the simultaneous binding of receptor and co-receptor is essential for the release of DNA into the host. For Phab24, we believe that a specific type of LOS molecule, which is not present in the mutant strain, is required to trigger the release of DNA into the bacterium, while the phage is able to bind via capsular polysaccharides in whose production *amsE* is involved. The work presented here points to phage resistance being caused by an alteration of the bacterial surface structure. Phage resistance caused by capsule loss has been described previously and the same publication also reports that a phage-escape mutant of a *A. baumannii* MDR strain showed a much reduced propensity to form biofilms (34). In our work, we also observed a decrease in the formation of biofilm when investigating the *amsE* gene knockout in the colistin sensitive reference strain ATCC17978.

Our study also showed that the two mutations that conferred phage-resistance result in decreased virulence in our *in vivo* model. Similar to our findings, it has been shown that phage escape mutants often show attenuated pathogenicity which, in some instances, is thought to be due to bacterial envelope modifications which are the basis of phage resistance (34, 36–39). Increasing evidence indicates that phage-resistance is a “trade-off” that results in decreased virulence and increased susceptibility to antibiotics (40). Therapeutic bacteriophages are commonly deployed together with antibiotics in clinical therapy. This is often done despite the fact that the bacteria display antibiotic resistance to the used compounds *in vitro*, due to the often observed phage-antibiotic synergy which we have also observed here (11, 28, 41–43). In our work, we saw a decrease in apparent MIC when colistin was used in combination with phage Phab24. The emerging phage-resistant bacterial clones often exhibited increased levels of antibiotic sensitivity. Previously, it has been proposed that phages could be used to re-sensitise bacterial pathogens to antibiotics (34, 44, 45). When we attempted to elucidate the underlying mechanisms, we found that phage resistant mutants with disruptions in *amsE* resulted in a drastic increase in colistin sensitivity, while the original mechanism for colistin resistance remained unaffected.

## Acknowledgements

We thank Mark Toleman (University of Cardiff) for critically reading the manuscript, to Nick Scott (University of Melbourne) for helpful discussions. We thank Belinda Loh who has obtained no financial compensation nor salary from the affiliated institution for her work during the last year.

SL, XH, YY and BL contributed to study conception and design. XW performed the experiments. XW, SL, BL analysed the data. SL supervised the study. SL and BL wrote the manuscript. All authors approved the final manuscript.

## Materials & Methods

Isolation, purification and the genome of phage Phab24 is described elsewhere (Manuscript submitted to *Genome announcement*). Genome accession number: MZ477002.

DNA genome sequencing and analysis. Genomic DNA was extracted using Bacterial genome DNA isolation kit (Biomed, China), sequenced by Illumina HiSeq (150-bp paired reads) and assembled using Unicycler version 0.4.8 (46). Breseq was used to identify single point mutations (20).

Gene knockout or replacement and complementation were performed as described previously: (47).

Determination of bacterial growth rates was performed as described previously (48) with the following modifications for experiments that included phages: An MOI of 5 of a high-titre preparation was added to the culture with a negligible dilution.

Transmission electron microscopy was performed as described previously: (49). Micrographs were obtained with a JEOL JEM1010 at 80 kV. Scanning Electron Microscopy as described here: (21).

Phage adsorption. Adsorption was measured indirectly by quantifying free phage in solution. Overnight bacterial culture was diluted in LB and bacteria at 8×10^9^/mL were incubated with Phab24 at 2×10^9^/mL at 4D for 20 minutes. Cells were pelleted by centrifugation, before the supernatant was serially diluted and non-adsorbed phages quantified by spot titre.

Laboratory evolution experiment: The soft-agar overlay technique was used to obtain phage resistant colonies which are purified three times by re-streaking, after co-incubation of ATCC17978 (or XH198) with Phab24 (MOI = 1) for 3 hours at 37°C.

Colistin MIC was determined by the broth-dilution method as described previously: (14).

Lipid A isolation and structural characterisation: Lipid A isolation was performed as previously described (50) which was used for MALDI-MS.

Surface polysaccharide extraction were purified by hot aqueous phenol extraction according to a previously described protocol (51).

Biofilm assays were preformed as described previously: (52). Each sample was done five times, with 3 independent experiments.

Crystal Violet (CV) retention assay of planktonic cells: Overnight cultures were diluted in LB media to OD600 = 1. 1 mL of diluted culture was centrifuged and cells were washed with PBS, then resuspended. CV was added (final 0.01% w/v), vortexed and incubated for 10 minutes. Cells were washed 3 times, before being destained with 2 mL of 95% ethanol for 10 minutes. Cell-free supernatant was then transferred into spectroscopic cuvettes and absorbance is measured at 540 nm.

*Galleria mellonella* infection model. Larvae survival assay was performed as previously described (53). Ten larvae per group.

Checkerboard Assay was performed as described previously: (54). 4 x10^4^ bacteria, with MOI: 5000, 500, 50, 5, 0.5, 0.05, 0.005.

## Supplemental Materials

**Supplemental Figure 1,2,3: Composition of the bacterial envelope**. (1) Mass spectra with m/z values from 180 to 750 of polysaccharides isolated from the wildtype strain ATCC17978, or the Phab24 resistant, genetically engineered (reconstructed) *amsE* gene mutant on ATCC17978. (2) Mass spectra with m/z values from 750 to 1500 of polysaccharides isolated from the wildtype strain ATCC17978, or the Phab24 resistant, genetically engineered (reconstructed) *amsE* gene mutant on ATCC17978. (3) Mass spectra with m/z values from 1500 to 2300 of polysaccharides isolated from the wildtype strain ATCC17978, or the Phab24 resistant, genetically engineered (reconstructed) *amsE* gene mutant on ATCC17978.

**Supplemental Figure 4: Surface structure of bacterial cell envelope.** Thin sections of bacterial cells visualised by TEM. Two pictures representative of cells were chosen for each cell type. *ΔamsE*: knock-out of *amsE* gene in the ATCC reference strain. *ΔlpsBSP:* knock-out of *lpsBSP* gene in the ATCC reference strain.

**Supplemental Figure 5: Membraneous structures and cell aggregation observed during experimental protocols using the ATCC17978 *amsE* KO mutant:** (Top) Biofilm Assay (Bottom) Capsular staining.

**Supplemental Table 1:** A selection of Phage-resistant isolates with gene mutations that are not responsible for phage resistance. The mutations conveying resistance are indicated (unless unknown). Last column (right): Negative outcome of wildtype gene complementations in trans indicate that resistance is not mediated by these genes. Mutations were either detected by whole genome sequencing and confirmed by PCR (strains R5, R10, R22, R23, R39, R70, R81, R83, R86, R115, R125, R130, R132, R134, R137), or screened for using gene-specific primers.

**Supplemental Table 2:** Colistin MIC values of phage resistant isolates (individual strains) derived from XH198 determined by micro broth dilution method.

