## Supplemental Table 2 for "Phage Resistance Mechanisms Increase Colistin Sensitivity in *Acinetobacter baumannii*"

| | Colistin MIC ug/ | P24 sensitiv | pmrB mutation | glyT mutation | lps mutation | amsE mutation | traL mutation |
|---|---|---|---|---|---|---|---|
| R588 | 0,25 | - | No verification | No verification | No verification | No verification | No verification |
| R515 | 0,5 | - | No verification | No verification | No verification | No verification | No verification |
| R567 | 0,5 | - | No verification | No verification | No verification | No verification | No verification |
| R573 | 0,5 | - | No verification | No verification | NO | No verification | No verification |
| R584 | 0,5 | - | No verification | No verification | No verification | No verification | No verification |
| R544 | 1 | - | No verification | No verification | No verification | No verification | No verification |
| R571 | 1 | - | No verification | No verification | No verification | No verification | No verification |
| R703 | 1 | - | No verification | No verification | No verification | No verification | No verification |
| R83 | 2 | - | p.Gly 315 Asp | NO | No verification | No PCR product of amsE | NO |
| R506 | 2 | - | No verification | No verification | No verification | No verification | No verification |
| R518 | 2 | - | No verification | No verification | p.Asn179 Ile fs Ter7 | NO | No verification |
| R536 | 2 | - | No verification | No verification | No verification | No verification | No verification |
| R545 | 2 | - | No verification | No verification | NO | No verification | No verification |
| R587 | 2 | - | No verification | No verification | p.Asn179 Ile fs Ter7 | NO | No verification |
| R701 | 2 | - | No verification | No verification | No verification | No verification | No verification |
| R130 | 4 | - | p.Gly 315 Asp | NO | No verification | transposase insertion | NO |
| R505 | 4 | - | No verification | No verification | NO | No verification | No verification |
| R523 | 4 | - | No verification | No verification | No verification | No verification | No verification |
| R527 | 4 | - | No verification | No verification | No verification | No verification | No verification |
| R558 | 4 | - | No verification | No verification | No verification | No verification | No verification |
| R580 | 4 | - | No verification | No verification | No verification | No verification | No verification |
| R125 | 8 | - | p.Gly 315 Asp | p. Met 150 Cys fs Ter22 | No verification | NO | NO |
| R134 | 8 | - | p.Gly 315 Asp | NO | No verification | transposase insertion | NO |
| R504 | 8 | - | p.Gly 315 Asp | No verification | No verification | No verification | No verification |
| R522 | 8 | - | No verification | No verification | NO | p.Gln234Ter1 | No verification |
| R524 | 8 | - | No verification | No verification | No verification | No verification | No verification |
| R533 | 8 | - | No verification | No verification | No verification | No verification | No verification |
| R577 | 8 | - | No verification | No verification | No verification | No verification | No verification |
| R596 | 8 | - | No verification | No verification | NO | No verification | No verification |
| R86 | 16 | - | p.Gly 315 Asp | NO | No verification | transposase insertion | NO |
| R115 | 16 | - | p.Gly 315 Asp | NO | No verification | transposase insertion | NO |
| R132 | 16 | - | p.Gly 315 Asp | p. Met 150 Cys fs Ter22 | No verification | transposase insertion | NO |
| R528 | 16 | - | p.Gly 315 Asp | No verification | No verification | No verification | No verification |
| R537 | 16 | - | No verification | No verification | No verification | No verification | No verification |
| R551 | 16 | - | No verification | No verification | NO | No verification | No verification |
| R564 | 16 | - | No verification | No verification | No verification | No verification | No verification |
| R569 | 16 | - | No verification | No verification | No verification | No verification | No verification |
| R572 | 16 | - | No verification | No verification | No verification | No verification | No verification |
| R589 | 16 | - | No verification | No verification | No verification | No verification | No verification |
| R590 | 16 | - | No verification | No verification | NO | No verification | No verification |
| R705 | 16 | - | No verification | No verification | No verification | No verification | No verification |
| R710 | 16 | - | No verification | No verification | No verification | No verification | No verification |

| | | | | | | | |
|---|---|---|---|---|---|---|---|
| R549 | 32 | - | No verification | No verification | NO | No verification | No verification |
| R552 | 32 | - | No verification | No verification | No verification | No verification | No verification |
| R556 | 32 | - | No verification | No verification | No verification | No verification | No verification |
| R565 | 32 | - | No verification | No verification | No verification | No verification | No verification |
| R570 | 32 | - | No verification | No verification | No verification | No verification | No verification |
| R575 | 32 | - | No verification | No verification | No verification | No verification | No verification |
| R576 | 32 | - | No verification | No verification | No verification | No verification | No verification |
| R578 | 32 | - | No verification | No verification | No verification | No verification | No verification |
| R81 | 64 | - | p.Gly 315 Asp | NO | No verification | transposase insertion | NO |
| R553 | 64 | - | No verification | No verification | No verification | No verification | No verification |
| R568 | 64 | - | No verification | No verification | No verification | No verification | No verification |
| R707 | 64 | - | No verification | No verification | No verification | No verification | No verification |
| R501 | >64 | - | p.Gly 315 Asp | No verification | NO | No verification | No verification |
| R502 | >64 | - | No verification | No verification | No verification | No verification | No verification |
| R509 | >64 | - | No verification | No verification | NO | No verification | No verification |
| R511 | >64 | - | No verification | No verification | No verification | No verification | No verification |
| R514 | >64 | - | No verification | No verification | NO | No verification | No verification |
| R516 | >64 | - | No verification | No verification | No verification | No verification | No verification |
| R519 | >64 | - | No verification | No verification | NO | No verification | No verification |
| R520 | >64 | - | No verification | No verification | No verification | No verification | No verification |
| R531 | >64 | - | No verification | No verification | No verification | No verification | No verification |
| R535 | >64 | - | No verification | No verification | No verification | No verification | No verification |
| R550 | >64 | - | No verification | No verification | No verification | No verification | No verification |
| R555 | >64 | - | No verification | No verification | No verification | No verification | No verification |
| R557 | >64 | - | No verification | No verification | No verification | No verification | No verification |
| R559 | >64 | - | No verification | No verification | No verification | No verification | No verification |
| R560 | >64 | - | No verification | No verification | No verification | No verification | No verification |
| R561 | >64 | - | No verification | No verification | No verification | No verification | No verification |
| R562 | >64 | - | No verification | No verification | No verification | No verification | No verification |
| R574 | >64 | - | No verification | No verification | No verification | No verification | No verification |
| R581 | >64 | - | No verification | No verification | No verification | No verification | No verification |
| R582 | >64 | - | No verification | No verification | No verification | No verification | No verification |
| R585 | >64 | - | No verification | No verification | No verification | No verification | No verification |
| R591 | >64 | - | No verification | No verification | No verification | No verification | No verification |
| R593 | >64 | - | No verification | No verification | No verification | No verification | No verification |
| R594 | >64 | - | No verification | No verification | NO | No verification | No verification |
| R702 | >64 | - | No verification | No verification | No verification | No verification | No verification |
| R704 | >64 | - | No verification | No verification | No verification | No verification | No verification |
| R706 | >64 | - | No verification | No verification | No verification | No verification | No verification |
| R708 | >64 | - | No verification | No verification | No verification | No verification | No verification |
| R709 | >64 | - | No verification | No verification | No verification | No verification | No verification |
| R711 | >64 | - | No verification | No verification | No verification | No verification | No verification |

**Minus** refers to resistance to P24
pmrB: AUO97-RS02735, Two-component system sensor histidine kinase
glyT: AUO97-RS06920, Glycosyltransferase
lps: AUO97-RS03485, LPS/ LOS biosynthesis
amsE: AUO97-RS06900, Amylovoran biosynthesis
traL: AUO970RS06930,Polysaccharide biosynthesis protein or Translocase

**No verification** refers to no PCR screen
**NO** refers to no related gene mutation via PCR screen
