## Supplemental Figures for "Phage Resistance Mechanisms Increase Colistin Sensitivity in *Acinetobacter baumannii*"

### Supplementary Figure 1

A

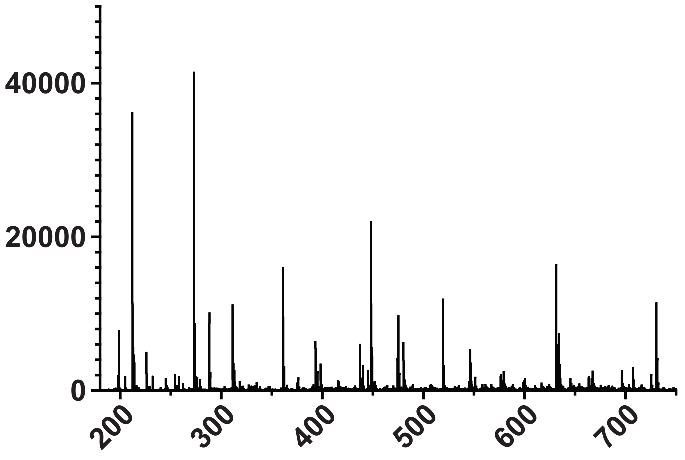


B

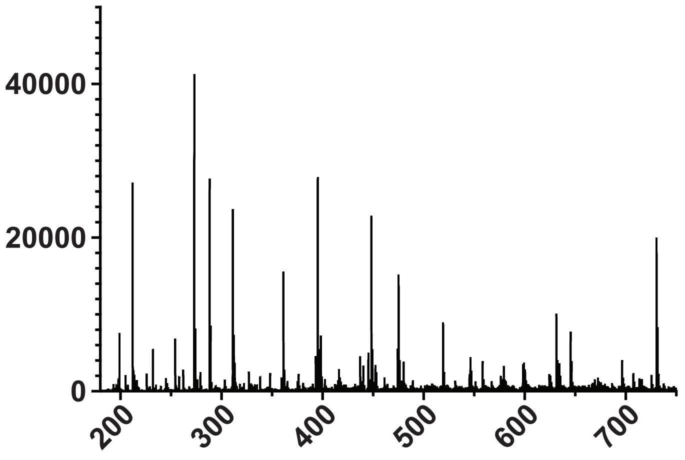

**Supplementary Figure 2**

A

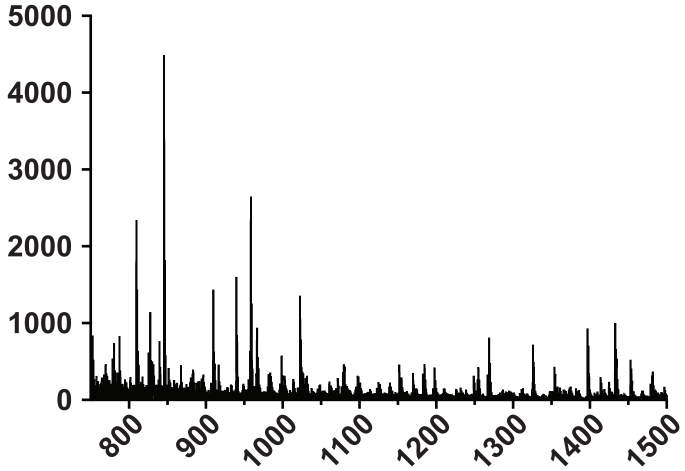


B

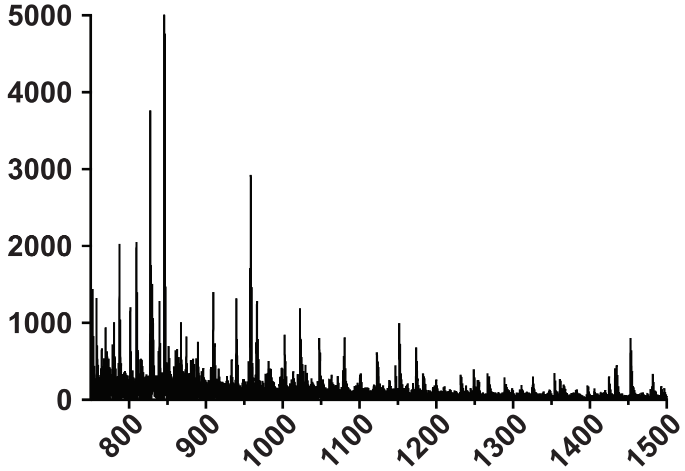

### Supplementary Figure 3

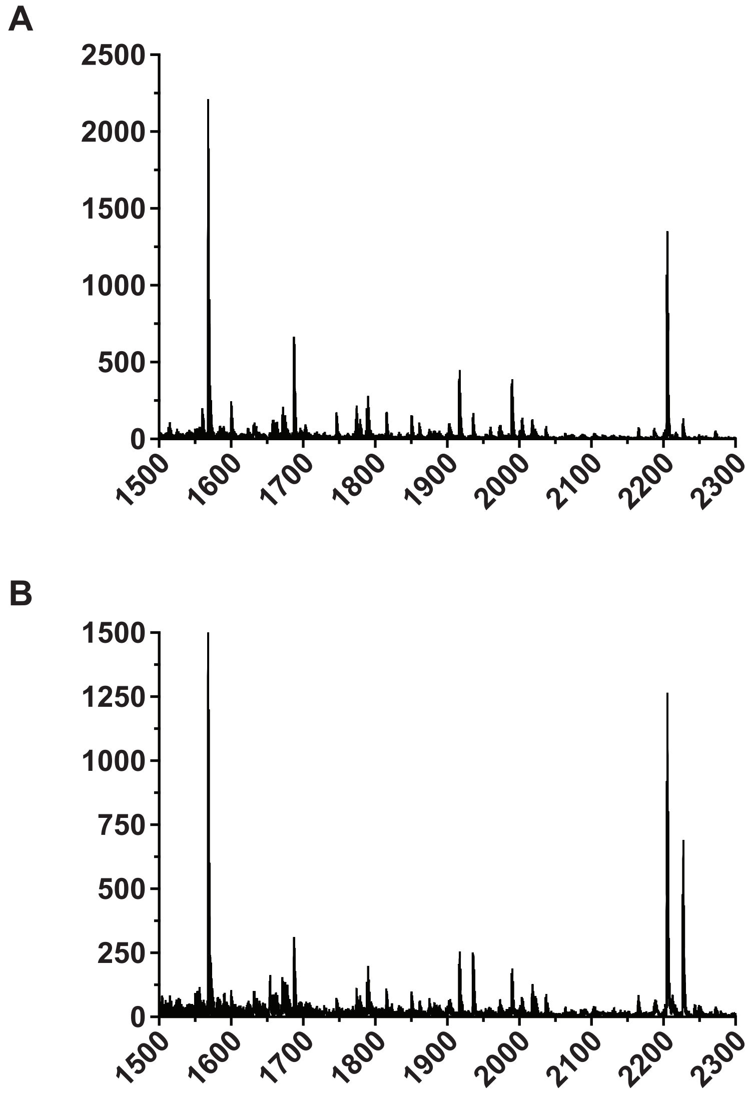

**Supplementary Figure 4**

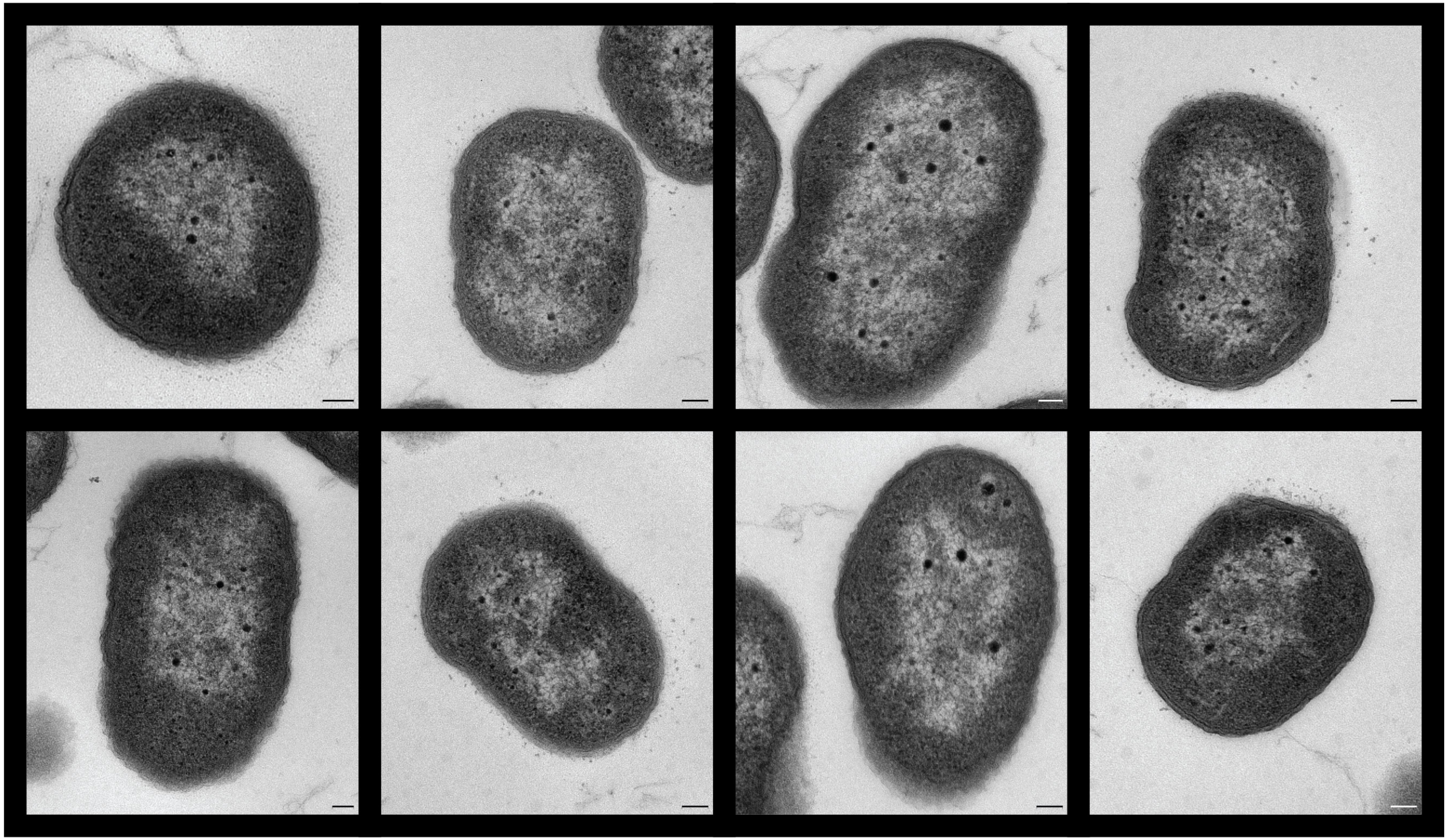

**XH198** **ATCC17978** **Δ*amsE*** **Δ*lpsBSP***

**Supplementary Figure 5**

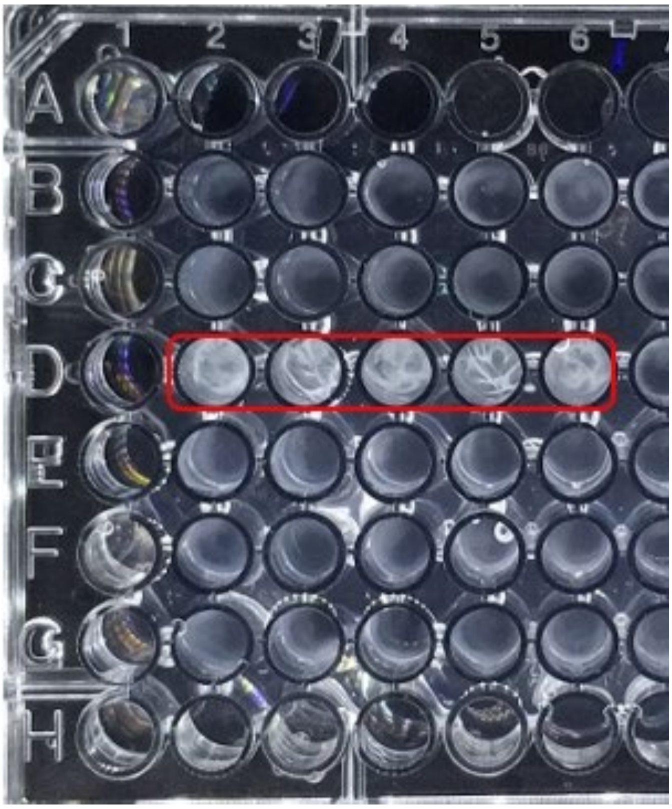

Biofilm assay in a 96 well plate. Cultures have been incubated for 72 hours. The amsE knockout mutant forms a kind of membraneous structure (wells D2-D6, red), which however does not retain the dye Crystal Violet that has been found to embed in biofilms.

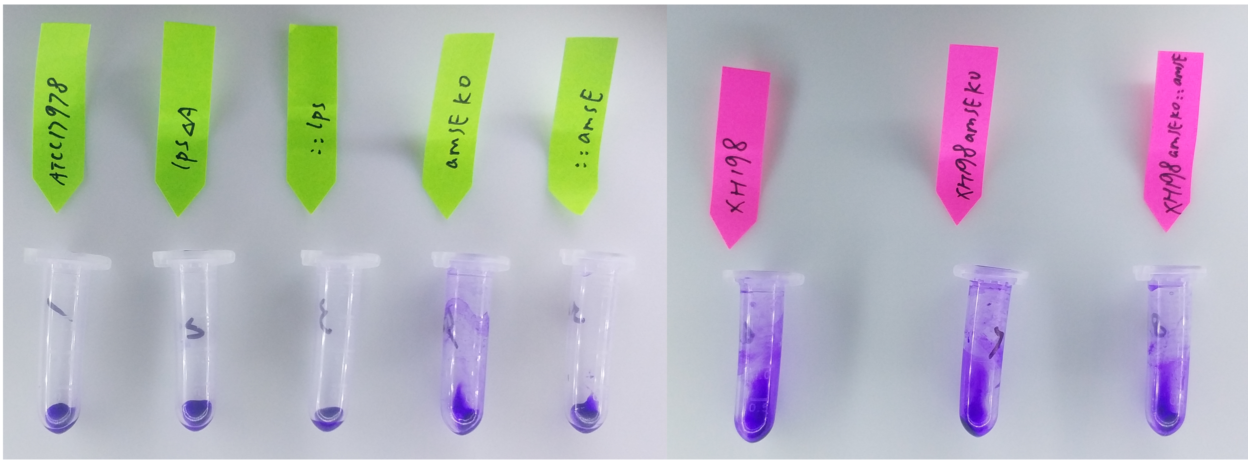


1 2 3 4 5 6 7 8

2 mL propylene sample tubes containing bacterial strains stained with the capsule staining dye Crystal Violet. The ATCC17978 *amsE* knockout mutant (4) is more adherent to the tube, similar to the XH198 strain, with or without the gene deletion (7) and complementation (8). The ATCC17978 complemented *amsE* knockout mutant (5) is similar to the reference strain (1) or the *lpsBSP* mutant (2) and the complemented *lpsBSP* mutant (3).
