## Supplemental Table 1 for "Phage Resistance Mechanisms Increase Colistin Sensitivity in *Acinetobacter baumannii*"

**SUPPLEMENTARY TABLE 1**

A selection of Phage-resistant isolates with gene mutations that are not responsible for phage resistance. The mutations conveying resistance are indicated (unless unknown). Last column (right): Negative outcome of wildtype gene complementations *in trans* indicate that resistance is not mediated by these genes. Mutations were either detected by whole genome sequencing and confirmed by PCR (strains R5, R10, R22, R23, R39, R70, R81, R83, R86, R115, R125, R130, R132, R134, R137), or screened for using gene-specific primers.

| Resistant Isolate | Gene mutation responsible for phage infection | Mutation | Putative gene function | Un-successful complementation with gene |
|---|---|---|---|---|
| R1 | *lpsBSP* | *abcT* (A1844T, p.Glu615Val) | ABC transporter | *abcT* |
| | | *actP* (A1144G, p.Thr382Ala) | acetate permease | *actP* |
| | | *phoH* (1011nt, 103th A loss) | phosphohydrolase | *phoH* |
| | | *decT* (753nt, G233A, p.Thr78Met) | di-trans,poly-cis-decaprenylcistransferase | NA |
| | | *dcaP* (1332nt, G395T, p.Pro132Glu) | DcaP-like protein | NA |
| | | *Udp* (720nt, T35C, p.Leu 12 Ser) | UDP-2,3-diacylglucosamine diphosphatase | NA |
| R2 | *lpsBSP* | *abcT* (A1844T, p.Glu615Val) | ABC transporter | *abcT* |
| | | *actP* (A1144G, p.Thr382Ala) | acetate permease | *actP* |
| | | *phoH* (1011nt, 103th A loss) | phosphohydrolase | *phoH* |
| | | *decT* (753nt, G233A, p.Thr78Met) | di-trans,poly-cis-decaprenylcistransferase | NA |
| | | *dcaP* (1332nt, G395T, p.Pro132Glu) | DcaP-like protein | NA |
| | | *Udp* (720nt, T35C, p.Leu 12 Ser) | UDP-2,3-diacylglucosamine diphosphatase | NA |
| R5 | *amsE* | *abcT* (A1844T, p.Glu615Val) | ABC transporter | *abcT* |
| | | *actP* (A1144G, p.Thr382Ala) | acetate permease | *actP* |
| | | *phoH* (1011nt, 103th A loss) | phosphohydrolase | *phoH* |
| | | *decT* (753nt, G233A, p.Thr78Met) | di-trans,poly-cis-decaprenylcistransferase | NA |

| Resistant Isolate | Gene mutation responsible for phage infection | Mutation | Putative gene function | Un-successful complementation with gene |
|---|---|---|---|---|
| | | *dcaP* (1332nt, G395T, p.Pro132Glu) | DcaP-like protein | NA |
| | | *Udp* (720nt, T35C, p.Leu 12 Ser) | UDP-2,3-diacylglucosamine diphosphatase | NA |
| R6 | Unknown | *abcT* (A1844T, p.Glu615Val) | ABC transporter | *abcT* |
| | | *actP* (A1144G, p.Thr382Ala) | an acetate permease | *actP* |
| | | *phoH* (1011nt, 103th A loss) | phosphohydrolase | *phoH* |
| | | *decT* (753nt, G233A, p.Thr78Met) | di-trans,poly-cis-decaprenylcistransferase | NA |
| | | *dcaP* (1332nt, G395T, p.Pro132Glu) | DcaP-like protein | NA |
| | | *Udp* (720nt, T35C, p.Leu 12 Ser) | UDP-2,3-diacylglucosamine diphosphatase | NA |
| R7 | *amsE* | *abcT* (A1844T, p.Glu615Val) | ABC transporter | *abcT* |
| | | *actP* (A1144G, p.Thr382Ala) | an acetate permease | *actP* |
| | | *phoH* (1011nt, 103th A loss) | phosphohydrolase | *phoH* |
| | | *decT* (753nt, G233A, p.Thr78Met) | di-trans,poly-cis-decaprenylcistransferase | NA |
| | | *dcaP* (1332nt, G395T, p.Pro132Glu) | DcaP-like protein | NA |
| | | *Udp* (720nt, T35C, p.Leu 12 Ser) | UDP-2,3-diacylglucosamine | NA |
| R8 | *lpsBSP* | *abcT* (A1844T, p.Glu615Val) | ABC transporter | *abcT* |
| | | *actP* (A1144G, p.Thr382Ala) | an acetate permease | *actP* |
| | | *phoH* (1011nt, 103th A loss) | phosphohydrolase | *phoH* |
| | | *decT* (753nt, G233A, p.Thr78Met) | di-trans,poly-cis-decaprenylcistransferase | NA |
| | | *dcaP* (1332nt, G395T, p.Pro132Glu) | DcaP-like protein | NA |

| Resistant Isolate | Gene mutation responsible for phage infection | Mutation | Putative gene function | Un-successful complementation with gene |
|---|---|---|---|---|
| | | *Udp* (720nt, T35C, p.Leu 12 Ser) | UDP-2,3-diacylglucosamine diphosphatase | NA |
| R10 | *amsE* | *abcT* (A1844T, p.Glu615Val) | ABC transporter | *abcT* |
| | | *actP* (A1144G, p.Thr382Ala) | an acetate permease | *actP* |
| | | *phoH* (1011nt, 103th A loss) | phosphohydrolase | NA |
| | | *decT* (753nt, G233A, p.Thr78Met) | di-trans,poly-cis-decaprenylcistransferase | NA |
| | | *dcaP* (1332nt, G395T, p.Pro132Glu) | DcaP-like protein | NA |
| | | *Udp* (720nt, T35C, p.Leu 12 Ser) | UDP-2,3-diacylglucosamine diphosphatase | NA |
| | | *isoS* (1698nt, 655th T loss, p.Ser 219 Pro fs Ter23) | 2-isopropylmalate synthase | NA |
| | | *LptD* (2439nt, 2395th G to T, p.Val 799 Phe) | LPS- assembly protein | NA |
| | | long-chain fatty acid--CoA ligase (1680nt,1240th T to A, p.Phe 414 Ile) | long-chain fatty acid--CoA ligase | NA |
| R12 | *lpsBSP* | *abcT* (1A1844T, p.Glu615Val) | ABC transporter | *abcT* |
| | | *actP* (A1144G, p.Thr382Ala) | an acetate permease | *actP* |
| | | *phoH* (1011nt, 103th A loss) | phosphohydrolase | *phoH* |
| | | *decT* (753nt, G233A, p.Thr78Met) | di-trans,poly-cis-decaprenylcistransferase | NA |
| | | *dcaP* (1332nt, G395T, p.Pro132Glu) | DcaP-like protein | NA |
| | | *Udp* (720nt, T35C, p.Leu 12 Ser) | UDP-2,3-diacylglucosamine diphosphatase | NA |
| R13 | Unknown, (not *amsE*) | *abcT* (A1844T, p.Glu615Val) | ABC transporter | *abcT* |
| | | *actP* (A1144G, p.Thr382Ala) | an acetate permease | *actP* |

| Resistant Isolate | Gene mutation responsible for phage infection | Mutation | Putative gene function | Un-successful complementation with gene |
|---|---|---|---|---|
| | | *phoH* (1011nt, 103th A loss) | phosphohydrolase | *phoH* |
| | | *decT* (753nt, G233A, p.Thr78Met) | di-trans,poly-cis-decaprenylcistransferase | NA |
| | | *dcaP* (1332nt, G395T, p.Pro132Glu) | DcaP-like protein | NA |
| | | *Udp* (720nt, T35C, p.Leu 12 Ser) | UDP-2,3-diacylglucosamine diphosphatase | NA |
| R125 | Unknown | *glyT* (448th A loss, p. Met 150 Cys fs Ter22) | Glycosyltransferase | *glyT* |
| R132 | Unknown, (not *amsE*) | *glyT* (448th A loss, p. Met 150 Cys fs Ter22) | Glycosyltransferase | *glyT* |

NA: No complementation attempted.

R1-R80 are Phab24 escape mutants from ATCC17978.
R81-R587 are Phab24 escape variants from XH198.
Based on the NCBI accession number CP018664.1, ATCC17978:

AUO97-07355, *abcT*, 1932nt

APP30310.1, *actP*, 1716nt

AUO97-03925, *phoH*, 1011nt.

AUO97-06920, *glyT*, 1164nt

AUO97-09800, 2-isopropylmalate synthase, 1698nt

AUO97-15295, LPS assembly protein *LptD*, 2439nt

AUO97-06215, long-chain fatty acid--CoA ligase, 1680nt

Interestingly, the phoH of the WT or ATCC17978 used for Sanger sequencing has 1012nt, while the phoH of CP018664.1 has 1011nt, the same as the mutated R variants.
